# *Atypus karschi* Dönitz, 1887 (Araneae: Atypidae): an Asian purse-web spider established in Pennsylvania, USA

**DOI:** 10.1101/2021.12.09.471987

**Authors:** Milan Řezáč, Steven Tessler, Petr Heneberg, Ivalú Macarena Ávila Herrera, Nela Gloríková, Martin Forman, Veronika Řezáčová, Jiří Král

## Abstract

The Mygalomorph spiders of the family Atypidae are among the most archaic spiders. The genus *Atypus* Latreille, 1804 occurs in Eurasia and northern Africa, with a single enigmatic species, *Atypus snetsingeri* Sarno, 1973, restricted to a small area in southeastern Pennsylvania in Eastern USA. This study was undertaken to learn more about genetics of that species, its habitat requirements and natural history. A close relationship to European species could be assumed based on *A. snetsingeri*’s occurrence on the eastern coast of the USA, however molecular markers (CO1 sequences) confirmed that *A. snetsingeri* is identical with *Atypus karschi* Dönitz, 1887 native to East Asia; it is an introduced species. The specific epithet *snetsingeri* is therefore relegated to a junior synonym of *A. karschi.* The karyotype of *A. karschi* has 42 chromosomes in females and 41 in males (X0 sex chromosome system). Chromosomes were metacentric except for one pair, which exhibited submetacentric morphology. In Pennsylvania the above-ground webs are usually vertical and attached to the base of bushes, trees, or walls, although some webs are oriented horizontally near the ground. It was found in a variety of habitats from forests to suburban shrubbery, and over a wide range of soil humidity and physical parameters. Prey include millipedes, snails, woodlice, carabid beetles and earthworms. The number of juveniles in excavated female webs ranged from 70 to 201. *Atypus karschi* is the first known case of an introduced purse-web spider. It is rarely noticed but well-established within its range in southeastern Pennsylvania.

## Introduction

Mygalomorph spiders of the family Atypidae are among the earliest divergent groups of spiders (1). They dig a burrow and construct a ‘purse-web’, usually in the form of a closed tube, that occupies the burrow and extends above the ground horizontally or vertically for prey capture. The webs are well-camouflaged with soil particles and plant debris and potential prey are sensed when they walk on the surface of the tube. The spider impales the prey through the silk with its long fangs and injects paralytic venom. It then makes a slit in the tube large enough to drag the prey inside, repairs the tear with new silk, and feeds on the prey (2–5). Atypid spiders spend their entire lifetime within their burrow in the silken web, 8–10 years for some females, and enlarge the burrow as they grow (6–8). Males abandon their burrows when they reach maturity and wander in search of females, and then mate within the female’s web. Egg-laying occurs within the maternal web and fully capable spiderlings emerge later. In contrast to most Mygalomorphs, atypid spiderlings utilize silk for aerial dispersal before establishing their first web (9,10). This ability may have allowed some species of atypids to colonize new areas, including those that were uninhabitable during the last glacial period (e.g., northern Europe, (11)).

There are currently three genera and 54 valid species of Atypidae (12). The genus *Atypus* Latreille, 1804 (34 species from Europe, Asia, North Africa and North America), spins an above-ground web that is tubular and typically lays horizontally and parallel to the soil surface. In *Sphodros* Walckenaer, 1835 (seven species from eastern North America) the above-ground web is tubular and usually attached vertically to trees and other vegetation. In the genus *Calommata* Lucas, 1837 (13 species from Africa and Asia) the above-ground web is a flat circular pouch set on the soil surface (13).

The center of diversity of the genus *Atypus*, based on the number of species, is in southeastern Asia and at least three species live in the western Palearctic. Despite the number and widespread distribution of *Atypus* species, they are secretive animals, and little is known about their habitat requirements, natural history, and genetic variation. In central Europe, particular *Atypus* species prefer sites with a microclimate regime resembling the climate of the glacial refuges from where they colonized the region (14). The species that live on open steppe habitats require soils rich in calcium that maintain a favorable air humidity in spider burrows. The species that do not require calcic soils occur only in habitats sheltered by woody vegetation, and their webs are hidden in detritus (15). As such, the European *Atypus* spiders are indicators of stable relic habitats and considered optimal flagship species in the conservation of disappearing relic xerothermic habitats (8).

Currently there are 16 Atypidae species with recorded DNA sequence data, eleven of which represent the genus *Atypus* (16). In contrast, only four species of Atypids, also in the genus *Atypus*, have been studied for their chromosomal constitution: *Atypus affinis* Eichwald, 1830; *Atypus karschi* Dönitz, 1887; *Atypus muralis* Bertkau, 1890; and *Atypus piceus*, Sulzer, 1776. The reported diploid number ranges from 14 to 44, and sex chromosome systems XY, X0, and X_1_X_2_0 have been described (17–19). There are no data on other chromosome features, such as constitutive heterochromatin or nucleolus organizing regions (NORs). Those chromosome markers have been sporadically examined in Mygalomorphae (17,20).

This study looked at the genetics and habitat requirements of the lone species of *Atypus* found in North America, *Atypus snetsingeri* Sarno, 1973 (21). This spider appears to be restricted to a small geographic area near Philadelphia, Pennsylvania in eastern USA (22). It is morphologically similar to *Atypus karschi* of Asia (7,23,24) hypothesized that it was probably introduced to North America by human activity. To help resolve its relationship with other Atypids, the karyotype and genetic barcode (CO1) were developed for *Atypus snetsingeri* to compare with other *Atypus* species, along with observations on habitat associations and natural history.

## Material and Methods

### Study locations

In November 2013 we visited eight sites in Delaware County, Pennsylvania, that were known to have *Atypus snetsingeri* populations (Tessler, personal observations). The sites ranged from semi-urban areas near the Type locality to wooded county parks along riparian corridors where purse-webs were common. The primary site used for detailed web observations, specimen excavation and collection was a fallow field surrounded by forest at the Tyler Arboretum (Media, PA). That field was mowed annually to control invasive plants and facilitated access to the webs.

### Habitat and natural history

At each site, we assessed the primary vegetation cover and soil type. The land orientation of the web location was measured using a compass and the slope angles using an optical reading clinometer to the nearest 0.5°. Soil penetration resistance was measured as described by Srba & Heneberg (25), where higher values reflect mechanical impedance for burrowing.

The range of web sizes (tube diameter) was visually assessed in the field and prey were noted by identifying remnants of invertebrates found attached to webs. The complete webs of 18 adult females were excavated on 5–9 November 2013. The length of the purse-webs were measured, distinguishing the below-ground and above-ground sections by coloration and attached soil. The size of the females was characterized by measuring the length of the carapace along the midline. When spiderlings were present, their number was counted. Voucher specimens from this study were deposited at the Crop Research Institute, Prague, Czechia.

### Statistical analysis

We used Pearson’s correlation test to analyze the correlation between carapace size and tube parameters (depth of the burrow, length of the capturing tube, and total length of the whole web) and to analyze the correlation between individual tube parameters. We evaluated the correlation between female size and number of offspring using the Spearman’s correlation test. The difference in body size between females with offspring and females without offspring was analyzed using the Welch two sample *t*-test. We tested the two variances in the subterranean and surface part of the tube by F-test. Normality was tested by Shapiro-Wilk normality test. Data were analyzed in the statistical software R 3.6.2 (26). The means are given with ± the standard error of the mean as a measure of sampling distribution.

### Karyotype analysis

Chromosome preparations were obtained from gonads of one immature male (testes present) and one mature female (ovary present). We followed the spreading technique described for mygalomorphs by Král et al. (20) except for fixation time (10 and 20 min). The standard preparations were stained by 5% Giemsa solution in Sörensen phosphate buffer for 25 min.

The evaluation of the karyotype was based on five mitotic metaphases. The chromosome measurements were carried out using ImageJ software (27). The relative chromosome lengths were calculated in each specimen independently as a percentage of the total chromosome length (TCL) of the haploid set, including sex chromosome. Chromosome morphology was classified according to Levan et al. (28).

Our study also includes detection of constitutive heterochromatin and nucleolus organizing regions. Male mitotic plates were used to visualize these markers. Constitutive heterochromatin was detected by C-banding following Král et al. (29). Preparations were stained by 5% Giemsa solution in Sörensen phosphate buffer for 75 min. NORs were visualized using fluorescence in situ hybridization (FISH) with a biotin-labeled probe for 18S rDNA sequences. The probe was obtained from *Dysdera erythrina* Walckenaer, 1802 (Dysderidae). FISH, probe detection by streptavidin-Cy3 and signal amplification was performed as described by Forman et al. (30).

### DNA extraction, amplification and sequencing

We isolated the DNA from legs of three *A. snetsingeri* individuals. We washed the ethanol-fixed legs twice for 15 min using 1 ml of 10 mM Tris-HCl (pH 7.5) with 5 mM EDTA. Subsequently, we extracted the DNA using a NucleoSpin Tissue XS kit (Macherey-Nagel, Düren, Germany) according to the manufacturer’s instructions. We then amplified the DNA using primers targeting nuclear (ITS2) and mitochondrial (CO1) loci using the following polymerase chain reaction mix: 10 mM Tris-HCl (pH 8.8), 50 mM KCl, 1.5 mM MgCl_2_, 0.1% Triton X-100, 0.2 mM dNTP (each), 1 μM forward primer, 1 μM reverse primer, 0.5 U of Taq DNA polymerase (Top-Bio, Prague, Czech Republic), and 300 ng of extracted genomic DNA. The total reaction volume was 25 μl. To amplify the ITS2 locus, we used the primers ApicITS2FW2 (5’-CGATGAAGAACGCAGCCAGCTGCGAG-3’; (31)) and RITS (5’-TCCTCCGCTTATTGATATGC-3’; (32)). To amplify the CO1 locus, we used the primers LCO1490 (5’-GGTCAACAAATCATAAAGATATTGG-3’; ((33)) and C1-N-2194 (5’-CTTCTGGATGACCAAAAAATC-3’; (34)). We performed the reaction using an Eppendorf Mastercycler Pro thermal cycler (Eppendorf, Hamburg, Germany) for 36 cycles with 15-s denaturation at 94 °C, 2-min annealing at 43–57 °C, followed by a 1–3-min extension at 72 °C. We initiated the cycling with a 2-min denaturation at 94 °C and terminated it after 5-min incubation at 72 °C. Subsequently, we purified the amplified DNA using USB Exo-SAP-IT (Affymetrix, Santa Clara, CA) and bidirectionally sequenced the amplicons using an ABI 3130 DNA Analyzer (Applied Biosystems, Foster City, CA). For the three individuals of *A. snetsingeri* analyzed in their ITS2 locus and two for their CO1 locus, all the obtained ITS2 and CO1 sequences were identical. The resulting consensus DNA sequences were submitted to NCBI GenBank under accession numbers MT957000-MT957001 (CO1) and MT957146-MT957148 (ITS2).

### Alignments and phylogenetic analyses

We aligned the newly generated sequences with those of nine *Atypus* spp. obtained from NCBI GenBank as of September 7, 2020, and sequences of the corresponding outgroups by using MUSCLE (35,36) (gap opening penalty −400, gap extension penalty 0, clustering method UPGMB, lambda 24). We manually corrected the alignments for any inconsistencies, trimmed the aligned sequences to ensure that they all represent the same extent of the analyzed locus, removed short-length sequences from the alignments, and used only trimmed sequences for further analyses. The trimmed ITS2 locus [containing partial 5.8S ribosomal DNA and partial (close to full-length) ITS2 sequences] corresponded to nt 62-385 (324 bp) of *Atypus baotianmanensis* Hu, 1994 KP208877.1. The trimmed CO1 locus (partial CO1 coding sequence) corresponded to nt 23-595 (573 bp) of *Atypus piceus* KX536935.1. For each locus, we calculated the maximum likelihood fits of 24 nucleotide substitution models. We used a bootstrap procedure at 1,000 replicates and the nearest-neighbor-interchange as the maximum likelihood heuristic method to determine the tree inference when the initial tree was formed using a neighbor joining algorithm. We used best-fit models for the maximum likelihood phylogenetic analyses, including the estimates of evolutionary divergence between sequences.

## Results

### Phylogenetic analysis

Analysis of the DNA of *A. snetsingeri* has clarified its identity and the unusual presence of the genus in North America. We found that the CO1 locus (Fig. 6A) had a 100% sequence similarity (genetic distance of zero) with the matching 639bp-long CO1 locus of *A. karschi* (SDSU_MY4706) from the Honshu island, Japan (Hedin et al. 2019). After *A. karschi* the most closely related species for which sequences were available was the Asian *Atypus heterothecus* Zhang, 1985, with a genetic distance of 0.131 ± 0.021 of base substitutions per site between sequences. The European species, *Atypus piceus* and *Atypus affinis*, were basal to *A. snetsingeri* as well as to the whole group of hitherto sequenced Asian *Atypus* spp. (Fig. 6A). Concerning the ITS2 locus, the sequences of only two other *Atypus* spp. are known (Fig. 6B); therefore, this hypervariable locus awaits future analyses when more comparative data are available. The genetic distance to the closest species already sequenced in the ITS2 locus, *Atypus baotianmanensis*, was 0.109 ± 0.022 of base substitutions per site between the sequences.

### Taxonomy

Based on an exact match of the genetic CO1 barcode data, the *Atypus snetsingeri* purse-web spiders in Pennsylvania appear to represent an introduced local population of the Asian species *Atypus karschi*. In the remainder of this paper those spiders are referred to as *Atypus karschi ‘*from Pennsylvania’. The specific epithet *snetsingeri* is relegated to a junior synonym of *karschi.*

#### *Atypus karschi* Dönitz, 1887

*Atypus snetsingeri* Sarno, 1973: Sarno 1973 (21): page 38, figs 1–9 (description of both sexes).

**Figure 1.**
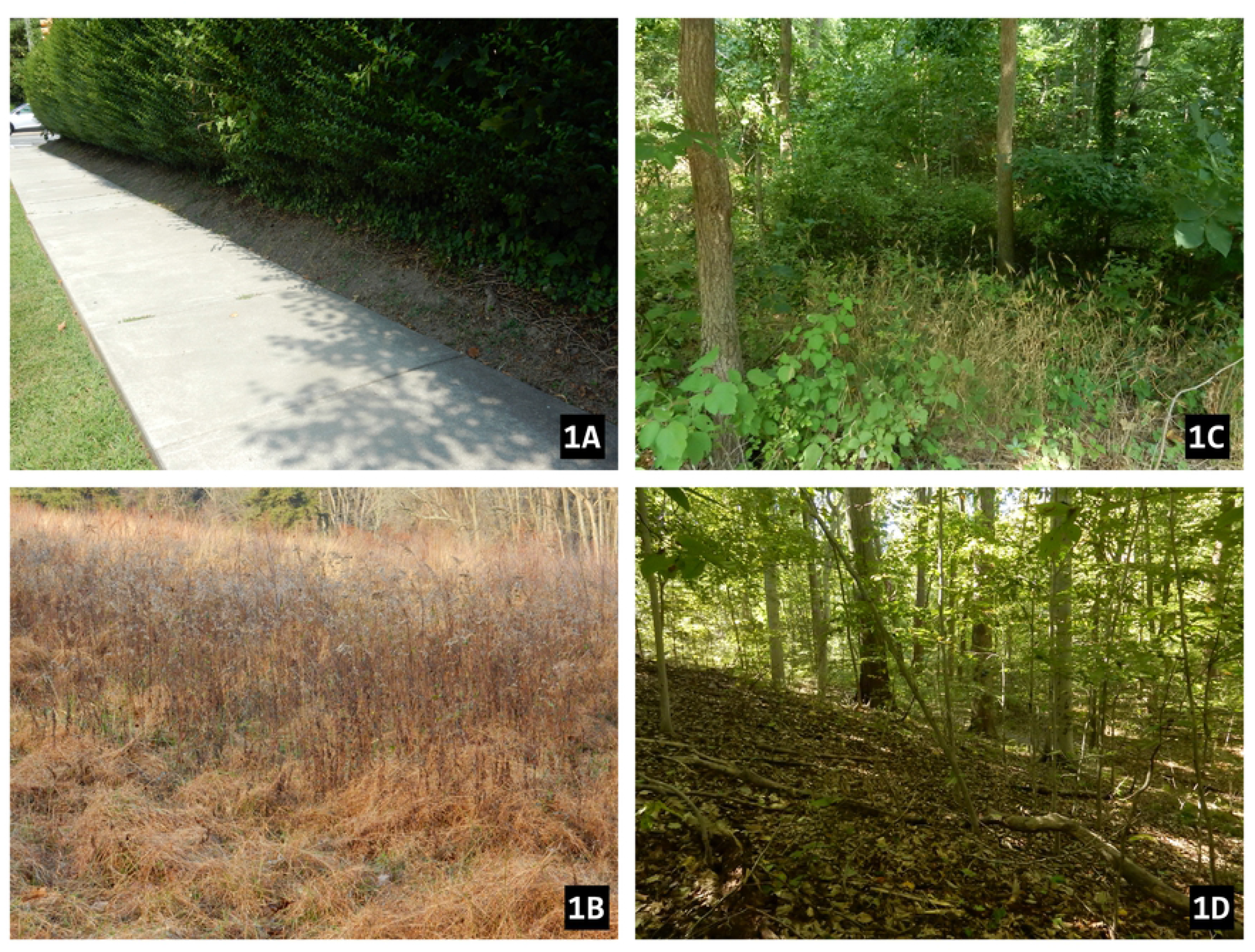
Habitats of *Atypus karschi* in Pennsylvania, USA; (1A) suburban bushes along Ellis Ave (~200 m from type locality of *Atypus snetsingeri),* (1B) fallow field at Tyler Arboretum, (1C) riparian woods, Swedish Cabin site on Darby Creek, (1D) forest, Smedley Park.

#### New synonymy

*A. snetsingeri* Gertsch and Platnick 1980 (23): page 11, figs 9, 13–20 (both sexes).

*A. snetsingeri* Schwendinger 1990 (7): page 360, fig. 18 (female).

##### Remarks

The synonymy was based on finding that the CO1 gene, used as a molecular barcode, of *snetsingeri* specimens from Pennsylvania was identical with that of *Atypus karschi* from the Honshu island, Japan (37).

### Karyotype

The female karyotype of *A. karschi* from Pennsylvania showed 2n = 42 chromosomes and the male had 2n = 41 (Fig. 5A), suggesting an X0 sex chromosome system. Chromosomes were metacentric except for one pair, which exhibited submetacentric morphology (Fig. 5B). The chromosome pairs gradually decreased in size, with the length of chromosome pairs in the male ranging from 7.13% to 3.31% of TCL and in the female from 6.13% to 3.31% of TCL. The sex chromosome was a metacentric element of medium size in both male (TCL = 4.27%) and female (TCL = 4.09%) (Fig. 5A, B). Concerning meiosis, pachytene nuclei were found in both the male and female specimen. In the male pachytene, the univalent X chromosome was on the periphery of the nuclei. X chromosome arms were often associated with each other during this period. Moreover, the X chromosome showed positive heteropycnosis (i.e., it was stained more intensively than other chromosomes). The other bivalents exhibited prominent knobs (Fig. 5C).

C-banded chromosomes exhibited small intercalar and terminal blocks of heterochromatin. The submetacentric pair showed a prominent large block at the terminal part of the long arm (Fig. 5D). It occupied on average 36% of the chromosome length (n = 10). The karyotype contained one NOR locus that was localized in the end of the long arm of the submetacentric pair (Fig. 5E). The NOR colocalised with the large block of heterochromatin and was of considerable size (37.2% of the chromosome length, n = 8).

### Habitat

The eight *Atypus karschi* sites that we visited in Delaware County in 2013 represented suburban neighborhoods, small wooded parks, narrow riparian zones along developed stream corridors, and protected parklands (Appendix 1). The purse-webs were located in a variety of habitats at those sites, including the shrubs along suburban sidewalks, slopes and bottoms of wooded valleys, beech forests and a fallow field that is mowed annually. Typical habitats of *A. karschi* in Pennsylvania are shown in Fig. 1.

The inclination (slope) of the sites varied from 0–40°, ranging from a flat field to riparian hillsides. Where a site in our study had a slope it usually faced the south but the azimuth of orientation varied from 95–340°, excluding only the coldest north and north-east exposures. The soil on slopes was usually not aggregated, was sandy or powdery, and of yellow or grey color below the shallow humus layer. In valley bottoms, the spider lived in fluvisol and in the suburbs in anthropogenic soils. Soil penetrability ranges from 0.5 to 3.25 (n = 14, mean 2.02 ± 0.31). The webs are typically associated with woody vegetation, and bush/shrub cover ranged from 5–100 % (mean 40 %) and tree cover from 0–90 % (mean 50 %). The soil surface where webs occurred was without moss, and the herbaceous cover was usually sparse (from 0–90 %, mean 20 %).

### Natural history

The above-ground webs we observed were vertical and mostly attached to the base of thin stems of bushes or on trees (Figs 2, 3), but a few were attached to rock. In early November three size categories were visually distinguished in the field by their relative web diameters, representing small and medium juveniles, and adult females. According to prey remnants found on their webs, they feed on millipedes (*Julida* and *Polydesmus* sp.), snails (*Cochlicopa* sp.), woodlice *(Porcellio* sp.) and carabid beetles.

**Figure 2.**
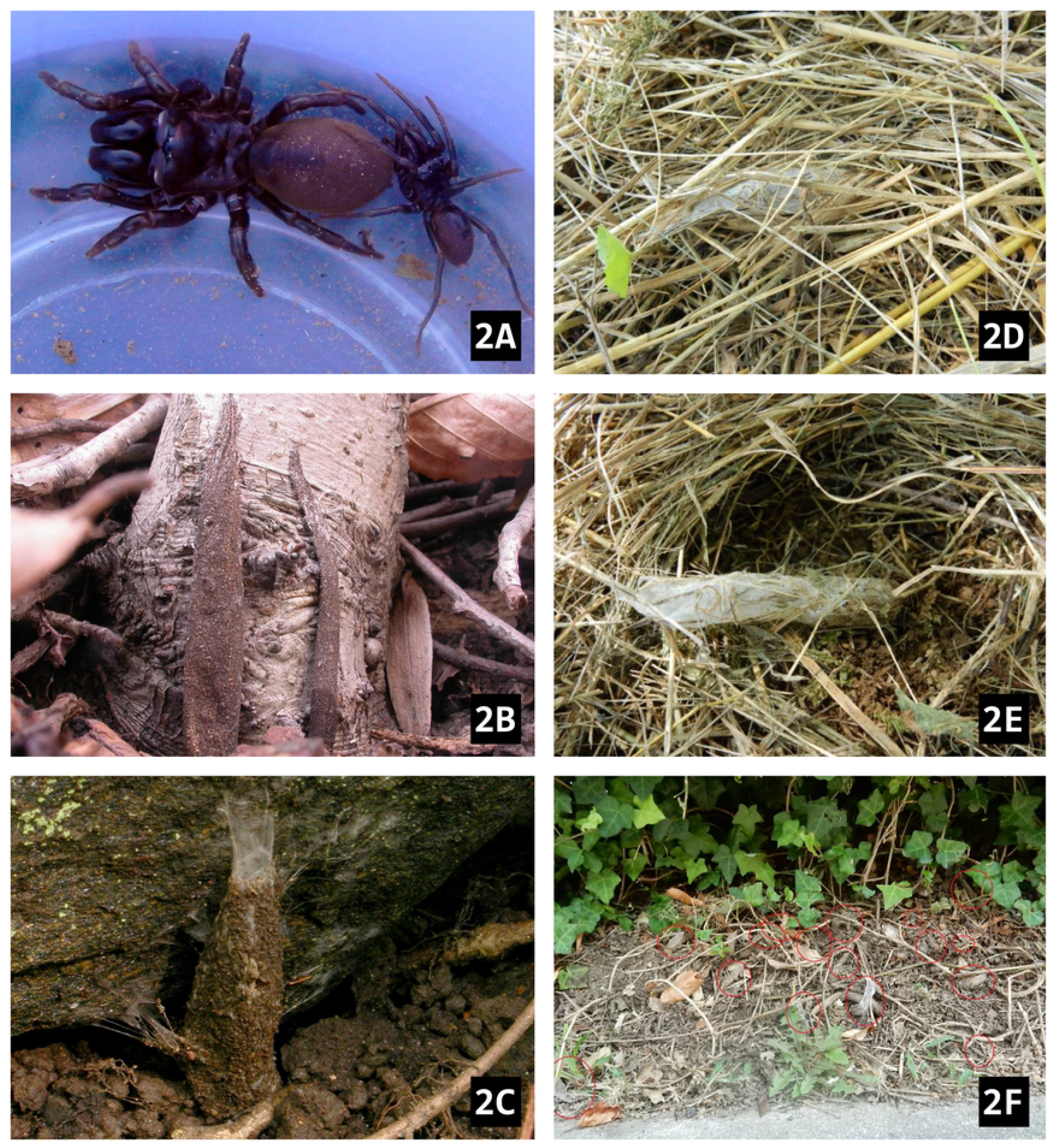
*Atypus karschi* and its webs in Pennsylvania, USA, (2A) adult female and male (on the right), (2B) vertical web attached to the base of a tree and (2C) to a boulder, (2D) horizontal web covered in thatch, (2E) thatch removed, (2F) view of trimmed ground in front of bushes with 16 purse webs.

**Figure 3.**
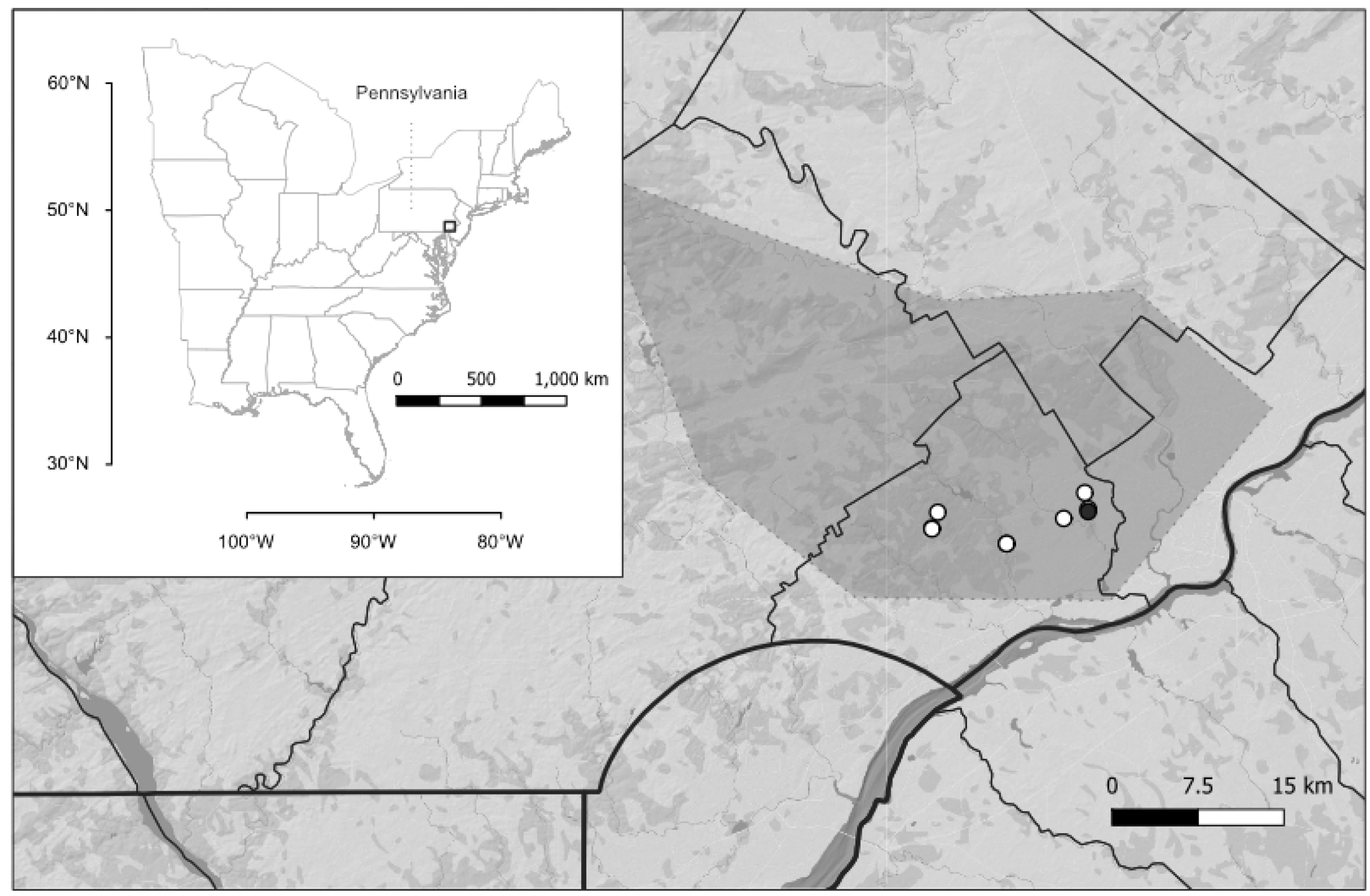
The distribution of *Atypus karschi* in Pennsylvania, USA. The peaks of the polygon represent the outermost sites. The circles mark the sites described in this study. The black circle is the type locality of *Atypus snetsingeri*.

Eleven out of 18 adult female webs that were excavated contained juveniles. There was no significant difference between the body size (carapace length) of the females with (n = 11) and without (n = 7) juveniles (all females: n = 18, min 5.04 mm, max 6.18 mm, mean 5.68 ± 0.09 mm) (Welch two sample *t*-test t = 1.45, *p* = 0.17). The number of juveniles ranged from 70 to 201 (n = 10, mean 121.30 ± 11.66) and did not correlate with the body size of the female (Pearson’s correlation n = 10, r = 0.32, *p* = 0.37). The length of the subterranean section of tube associated with the burrow ranged from 6 to 10 cm (n = 13, mean 8.3 ± 0.3 cm) and did not correlate with the body size of the spider (Pearson’s correlation, n = 13, r = −0.44, *p* = 0.13). The length of the above-ground tube ranged from 5 to 13 cm (n = 14, mean 8.54 ± 0.67 cm) and also did not correlate with the body size of the spider (Pearson’s correlation, r = 0.02, *p* = 0.94). The length of the above-ground purse-web was more variable than the length of its subterranean part (F test, n = 13, F = 0.23, *p* = 0.018) (Fig. 4). The ratio of below-ground/above-ground lengths ranged from 0.62 to 1.80 (n = 14, mean 1.09 ± 0.08) and did not correlate with the body size of the spider (Pearson’s correlation, r = 0.02, *p* = 0.94).

**Fig. 4.**
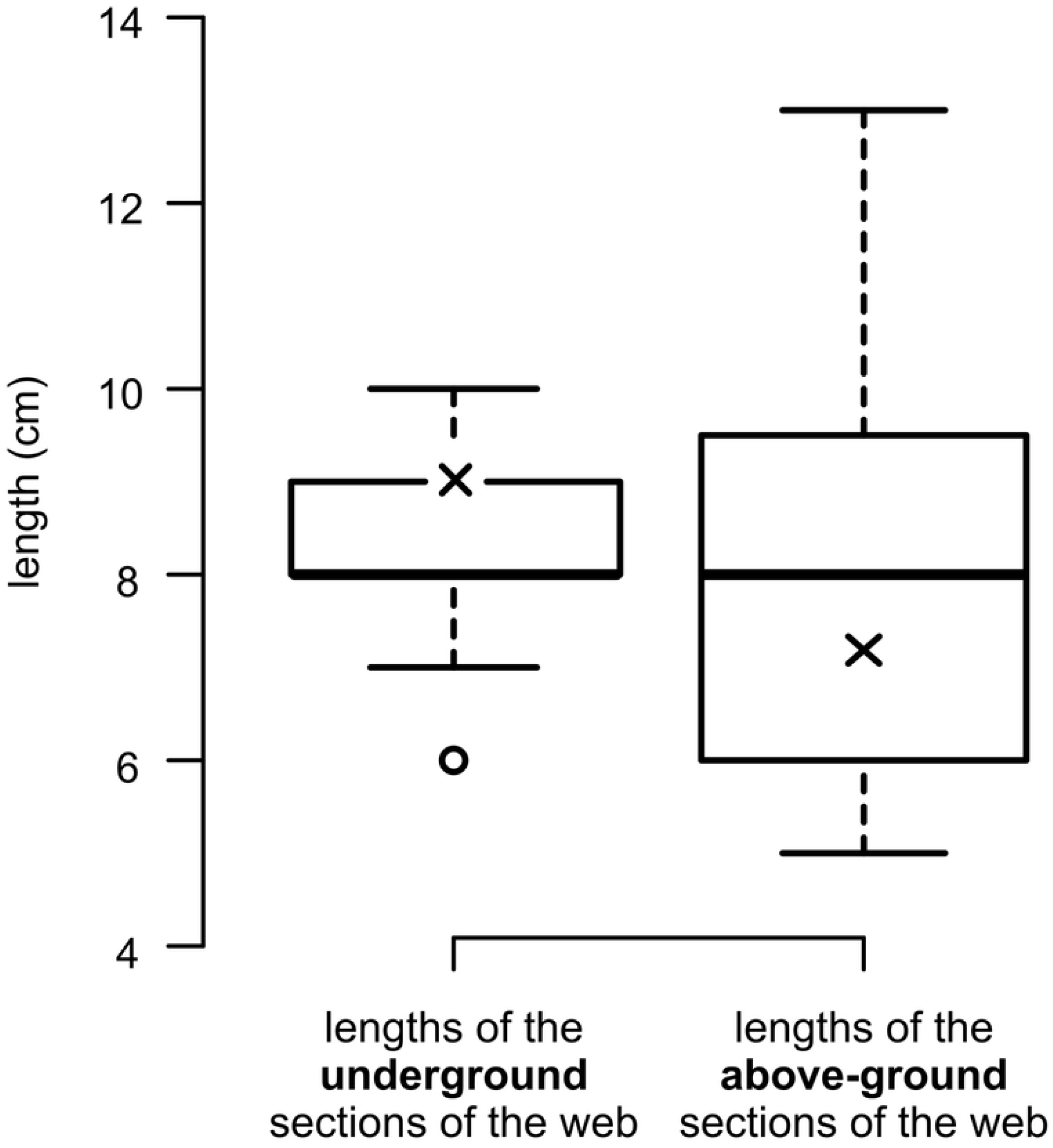
Boxplots showing the variation of the below-ground and above-ground lengths of excavated purse-webs of *Atypus karschi,* Pennsylvania, USA (n = 13). The means are indicated by an x and the hollow dot indicates an outlier (less than the 25th percentile minus 1.5 × Interquartile range).

**Figure 5.**
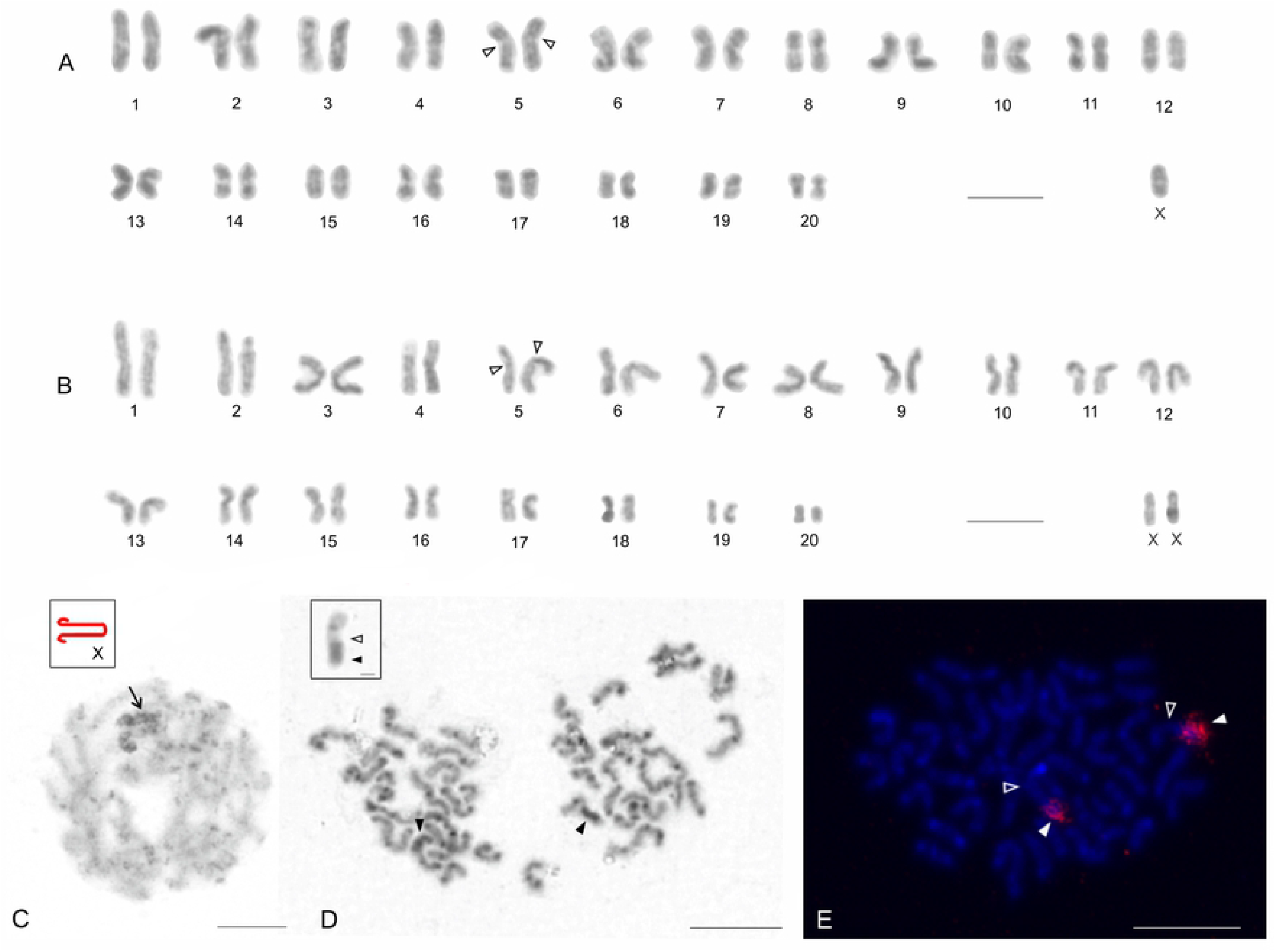
Chromosomes of *Atypus karschi,* Pennsylvania, USA. **A, B.** Male (A) and female karyotypes (B), stained by Giemsa, based on mitotic metaphase. 2n♂ = 41, X0; 2n♀ = 42, XX. Empty arrowhead – centromere of submetacentric pair. **C.** Male pachytene. Note heterochromatic X chromosome on the periphery of the nucleus and prominent knobs on the bivalents. Inset: scheme of sex chromosome. Note an association of X chromosome arms. Arrow – sex chromosome. **D.** Male mitotic metaphase, C-banding. Chromosomes exhibit intercalar and terminal heterochromatin blocks. Inset: magnified submetacentric chromosome containing a large block of heterochromatin (from another mitotic metaphase). Arrowhead – a large block of heterochromatin, empty arrowhead – centromere. **E.** Male mitotic metaphase, detection of rDNA cluster (FISH). Note chromosomes of a submetacentric pair containing a terminal rDNA cluster at long arm. Arrowhead – rDNA cluster, empty arrowhead – centromere. Scale bars: 10 μm.

**Figure 6.**
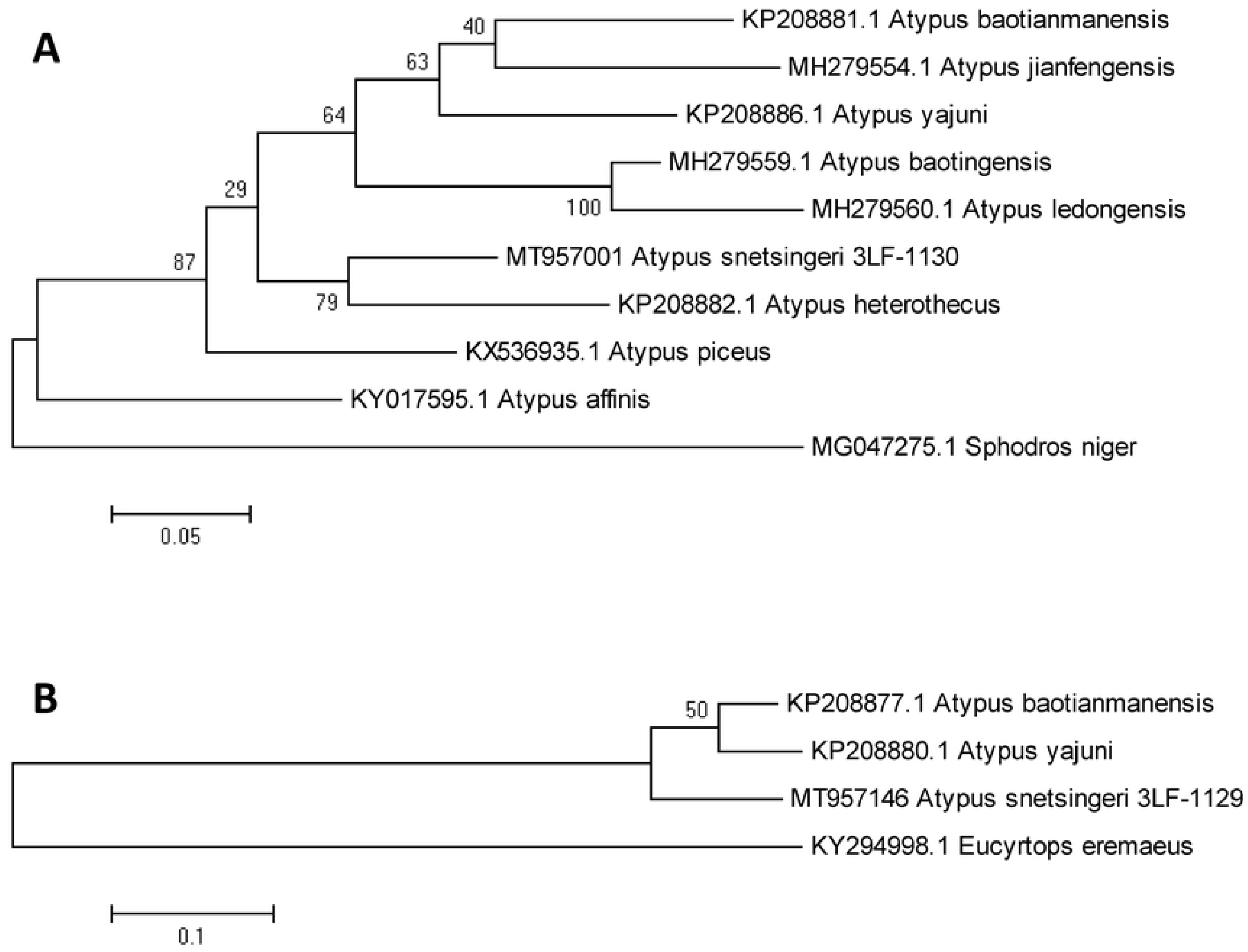
Phylogenetic analyses of the position of *Atypus karschi* (Pennsylvania, USA) in the genus *Atypus* based on the sequences of the CO1 (**A**) and ITS2 (**B**) loci by the maximum likelihood approach. The evolutionary history was inferred using the Tamura-Nei model (**A**) or the Kimura 2-parameter model (**B**), both with a discrete Gamma distribution used to model evolutionary rate differences among sites. The models were selected based on the highest Bayesian information criterion scores of the maximum likelihood fits. The trees are drawn to scale, with branch lengths indicating the number of substitutions per site. All codon positions, including noncoding positions, were included; the analyses were based on 573 positions (**A**) or 345 positions (**B**).

## Discussion

### Genetic identity of Atypus snetsingeri

The presence of a geographically isolated population of an *Atypus* species in North America, where the native purse-web spiders are in the genus *Sphodros*, has been mildly controversial. Due to the species’ location on the eastern coast of the USA a close relationship with European *Atypus* species could have been expected. However, morphologically, *A. snetsingeri* was known to closely resemble the Asian *A. karschi* (7,23). Raven (24) hypothesized that the single *Atypus* species in the USA was introduced by man.

The newly obtained molecular data for *A. snetsingeri* have resolved those questions by showing that the Pennsylvania species is more closely related to Asian species of *Atypus* than to European species. More specifically, *A. snetsingeri* was a genetic match with sequence data for *A. karschi* from Japan, affirming that the species represents an East Asian introduction. Based on these data we propose a formal synonymy for *A. snetsingeri*, which now becomes a junior synonym of *Atypus karschi.* Differences reported for morphological features compared to *A. karschi* in Asia probably represent intraspecific variation given the small number of *A. snetsingeri* specimens actually examined by researchers (7,23).

Parts of the genome of *“Atypus snetsingeri”* (based on NCBI sequences DQ639853.1, DQ680323.1 and KY016940.1) were used in spider phylogeny studies to represent the genus *Atypus* (37–39) or the entire family Atypidae (40). Wheeler et al. (1) used *A. snetsingeri* and *A. affinis* data to represent *Atypus*, and added *Sphodros* for the family Atypidae. Recently the entire mitochondrial genome was sequenced for *Atypus karschi* in China (41), which is highly useful for further comparative studies of the Atypoidea.

### Karyology

Most karyotype data on spiders concerns araneomorphs (42), but some karyotypes of mygalomorph spiders have been published (17,18,20,43,44). Representatives of the superfamily Atypoidea display a similar range of diploid numbers as araneomorph spiders (from 14 to 47). Most Atypoidea also exhibit the X0 sex chromosome determination system, which may be the ancestral characteristic sex chromosome determination of this superfamily (20).

In the family Atypidae only four species in the genus *Atypus* have been studied cytogenetically. *Atypus karschi* in this study exhibits 2n♂ = 41, X0 and predominance of metacentric chromosomes, which is in accordance with the karyotypes of central European species *A*. *piceus* and *A. muralis* (18). These karyotype features could be ancestral within the genus *Atypus.* The karyotype of European *A. affinis* having 2n♂ = 14, XY, was derived from chromosomal complement 2n♂ = 41, X0 by series of chromosomal fusions leading to decreasing of diploid count and formation of a neo-sex chromosome system XY (18).

Remarkably, an earlier karyotype developed for *A. karschi* in Japan (19) differs considerably from those reported in this study from Pennsylvania. The karyotype reported from Japan consisted of approximately of 44 acrocentric chromosomes, including an X_1_X_2_0 system, not the 42 chromosomes and X0 pattern reported here. The discrepancy may be due to interpopulation variability, but although mygalomorph spiders exhibits considerable karyotype diversity (20), such an enormous degree of interpopulation karyotype variability is very unlikely. Therefore, we suggest that the karyotype data of the Japanese population may have been misinterpreted. The karyotype of *Atypus* is formed by relatively high number of small chromosomes, which makes it difficult to determine the precise diploid number and chromosome morphology. Moreover, the method of chromosome preparation used by Suzuki (19) did not include treatment with a hypotonic solution, so the spreading of chromosomes would have been less pronounced than in the present study using the methodology of Král et al. (20). Regarding determination of the sex chromosome system, a single metacentric X chromosome of an X0 system could be erroneously considered as two acrocentric X chromosomes of an X_1_X_2_0 system attached at one end during meiosis.

Within the framework of our cytogenetic analysis we were able to detect constitutive heterochromatin for the first time in the Atypoidea. Most chromosomes of *A*. *karschi* exhibited intercalar and terminal blocks of heterochromatin. The distribution of blocks suggests that 1) most intercalar blocks are placed at centromeric regions and 2) terminal blocks are formed at telomeric regions. This is consistent with the pattern of constitutive heterochromatin distribution most commonly found in spiders (43).

Nucleolus organizer regions (NORs) are chromosome domains comprised of tandemly repeated sequences of rRNA genes that aid formation of the nucleolus after division (45), and their location on chromosomes may have taxonomic value. These regions have been detected in ten species of mygalomorphs including four species of Atypoidea ((20), this study). The number of NORs in Atypoidea ranges from one to four loci, and they are always situated on chromosome pairs. NORs are usually detected by impregnation with silver nitrate or by fluorescent in situ hybridization (FISH) with rDNA probe, although the first technique can underestimate absolute number of NORs by visualizing only loci transcribed during previous cell cycle (46). However, most NOR detections in mygalomorphs have been performed by silver staining. Fluorescence in situ hybridization, which we applied to detect NORs of *A. karschi* in this study, have been used with only one other mygalomorph species, *Tliltocatl albopilosum* Valerio, 1980 (Theraphosidae) (20). Both species display one terminal NOR localized on a chromosome pair, which may be the ancestral condition for spiders (20). The NOR of *A. karschi* is associated with heterochromatin, which is a common feature of rDNA in eukaryotes (e.g., (47,48)). Comparison of the length of the rDNA cluster and heterochromatin block suggests that heterochromatin associated with the NOR is formed by inactivated rDNA. This pattern is in an agreement with the current model for NOR organization, in which major regions of rDNA are often inactivated and only a restricted fraction of rDNA is transcribed (49).

### Habitat

*Atypus karschi* in Pennsylvania appears to be undemanding regarding habitat requirements (see Appendix 1) and can be locally abundant where it occurs. The webs are built in soil of varied humidity and physical parameters and are associated with a variety of supports (trees, shrubs, grasses, rock, walls and fences) over a ground surface either covered by or nearly devoid of litter. Webs were found on flat terrain and on slopes of various inclinations and orientation. In Pennsylvania it occurs in wooded areas but is also reliably found in some suburban neighborhoods, where webs are built at the base of shrubs or along walls and fences. Miyashita (50) reported a very similar situation in Japan where *A. karschi* is “common” and “usually live(s) in shady and humid places such as woods and shrubberies.” Images posted on iNaturalist (51) of *A. karschi* in East Asia also support a tolerance of human-modified settings where they were encountered (wall, fence and stone garden).

In sharp contrast, European *Atypus* species usually require very specific edaphic conditions and are associated with specific vegetation types and sun-facing slopes (14). Unlike *A. karschi* in Pennsylvania, they are not found in habitats subject to recent or regular disturbance and are uncommon enough to be red-listed in all Central European countries. Their presence at a site is an indicator of a relic habitat worthy of conservation management (8,52).

### Natural history

The life history of *A. karschi* in Japan was studied in detail by Miyashita (50) under semi-outdoor conditions and reported with prior data from Aoki (53) and Yaginuma (54). Basic natural history parameters of *A. karschi* in eastern Asia and *A. snetsingeri* in the USA are contrasted in Table 1 and indicated a similarity in every respect (body size, ontogeny, phenology, fecundity, morphology of webs, environment). No difference was found that would refute the conspecificity of the Pennsylvania population with Asian *A. karschi*.

**Table 1.** Natural history parameters (body size, ontogeny, phenology, fecundity, morphology of webs, environment) reported for *Atypus karschi* in Japan and for the introduced population known as *A. snetsingeri* in Pennsylvania, USA.

| Natural history parameter | *Atypus karschi* - Japan | *Atypus snetsingeri* - USA |
| --- | --- | --- |
| **Body size** |  |  |
| Carapace length of males | 3.87–4.23 mm (55) | 3.2–4.6 mm (21) |
| Carapace length of females | 4.77–5.76 mm (55) | 3.4–7.0 mm (21) |
| **Ontogeny** |  |  |
| No. of eggs | mean 124, maximum 270<br>(50) | mean 121, maximum 201<br>(this study) |
| No. of moults before reaching maturity | 8–9 in males, 9–11 in females (50) | unknown |
| Age of maturation | 3 years (50) | possibly 3 years, based on three concurrent web size categories in the population (this study) |
| **Phenology** |  |  |
| Mating season | June – (July) August (50,53) | June–August (21,22) |
| eggs | July (August) - September (50,53,54) | July-September (22) |
| larvae | October (56,57) | September (22) |
| 1st nymphal instar | late October–April (dispersion) (50,53,53) | September–March (dispersion) (22) |
| **Morphology of webs** |  |  |
| Orientation of the capturing tube | vertically attached to the tree trunk or rock (57) | vertically attached to the tree, hedge or wall (21) or horizontally oriented in grass and thatch (22) |
| Length of the capturing tube | Up to 20 cm (almost the same as the depth of the burrow) (57) | Up to 25 cm (21) |
| Depth of the burrow | Up to 20 cm (50,57) | Up to 20 cm (21) |
| **Environment** |  |  |
| Microclimate | Shady and moist, in the forest close to the trees, rocks or bamboo (50,57) | Mostly shady and moist, in litter and areas with loose soil (this study) |
| Habitat | Forests and shrubs (50,57) | Forests and shrubs, disturbed areas (this study) |

Although Gertsch and Platnick (23) contemplated whether the above-ground purse-web orientation could be useful to distinguish between *Atypus* (horizontal tubes) and *Sphodros* (vertical tubes), species in both genera can and do make both kinds of webs (58). In our study we observed only vertical webs of *A. karschi* at the sites visited, but the spiders are known to make horizontal webs in thatch and grass (22). In Tyler’s fallow field, for example, vertical webs can be found on plant stems within a few centimeters of horizontal webs in surrounding grasses. While vertical tubes are characteristic of North American *Sphodros* species, *Sphodros niger* Hentz, 1842 may preferentially build horizontal tubes, at least in some settings (59,60). Mckenna-Foster et al. (61) found that *Sphodros rufipes* Latreille, 1829 in New England will use whatever support is available and many webs were close to the ground. The suggestion that horizontal webs are an adaptation to prey capture under the snow (7) may ignore the function of vertical webs at ground level. In Pennsylvania *A. karschi* habitats experience snow and cold temperatures each year. In Tyler’s field the horizontal tubes laying near the soil surface tend to be well-buffered by leaf litter or thatch, but basal sections of vertical webs are similarly buffered and may likewise function normally in a subnivean environment when both prey and spiders are active (Tessler, personal observations).

In this study we measured the webs of fourteen adult females from a fallow field with homogeneous soil. We found the overall length of the webs were shorter than those observed by Sarno (21) around a house foundation and on shrubs in a suburb (see Table 1), probably reflecting different conditions and prey availability between sites. The length of the aerial tube was more variable than that of the underground part (Fig. 4). Less variation in the underground web length may reflect a minimum depth of the burrow necessary for suitable microclimate, constraints imposed on digging, or the shallow soil frost depth in winter. Depth of burrows differs among European *Atypus* species, where the species living in arid habitats tend to dig deeper burrows than those living in woody vegetation (8).

Concerning the number of juveniles found within maternal webs, *A. karschi* in Pennsylvania (max. 201 juveniles) has a similar number as *A. karschi* in Asia. Likewise, European *Atypus* species also have large broods (*A. affinis* max. 191, *A. piceus* max. 168, *A. muralis* max. 150; M. Řezáč, personal observations).

Prey we observed for *A. karschi* in Pennsylvania were mostly ground-based invertebrates and favored millipedes, similar to observations on *S. niger* in New England (60). *Atypus karschi* in Pennsylvania has also been observed feeding on earthworms, and will readily capture orthopteroids and other insects that contact the web while climbing vegetation, including the pestiferous spotted lanternfly (Hemiptera: Fulgoridae: *Lycorma delicatula* White, 1845) that was recently introduced into Pennsylvania from Asia (Tessler, personal observations).

### *Range of* Atypus karschi *in Pennsylvania*

*Atypus karschi* seems to possess several preadaptations that allowed it to successfully colonize southeastern Pennsylvania following its introduction. First, it occurs over a wide area in eastern Asia with a similar climate (Japan (57); Chinese provinces Hebei, Anhui, Sichuan, Guizhou, Hubei, Hunan, Fujian (55); Taiwan (62); Korea’s Ungil Mountain (63)). Second, it produces a large number of lightweight juveniles that are capable of aerial dispersal (22,50,64). Third, the species in Pennsylvania is ecologically plastic and does not appear to have specific edaphic or microclimatic requirements, even thriving in settings frequently impacted by humans.

The original description and first review of the species *A. snetsingeri* in Pennsylvania was based on specimens taken from two nearby suburban sites in Lansdowne and Upper Darby in eastern Delaware County near Philadelphia (21,23). At that time, it was known be common and unnoticed in the surrounding areas within the Cobbs Creek and Darby Creek drainage basins (Tessler, personal observations). It has subsequently been sought and found in many (not all) of the forested riparian zones and wooded parklands across Delaware County and also at sites in adjacent areas of Philadelphia, Montgomery, Chester and Berks counties (Fig. Map). Many neighboring areas, including most of urban Philadelphia, remain unexplored (22). A few of those species determinations were based finding males, but the majority involved excavating a web to extract the spider and examine the sternum sigilla pattern and the posterior lateral spinnerets (PLS) to distinguish it from *Sphodros* species (23). In particular, *A. snetsingeri* has a distinctly four-segmented PLS, whereas the northern *Sphodros* species have only three segments (*S. niger, S. rufipes, S. atlanticus*).

Spiderlings of *A. karschi* in Pennsylvania use silk for aerial dispersal in the spring (22), which may have contributed to expanding its range from an original introduction locus. However, the association of these spiders with highly developed land and disturbed habitats suggests a wider transport opportunity via trees, woody shrubs and mulch moved within the region by landscaping and nursery industries (Tessler, personal observations).

Interestingly, *Sphodros* purse-web spiders are also found in Pennsylvania and adjacent states (23), but none have ever been reported in the same areas as *A. snetsingeri.* This is unsurprising because atypids and their webs are rarely noticed or reported even when they are locally abundant (22,65). Sightings of wandering *Sphodros* males reported in iNaturalist (66) indicate that *S. niger* is found in Pennsylvania west and north of the *A. karschi* area and southward in neighboring states, and that *S. rufipes* occurs in Maryland and New Jersey south and east of the Philadelphia area and northward into coastal New England. While perhaps provocative, those observations are not evidence of displacement of any local species by the introduction of *A. karschi.*

It is unlikely that the source and timing of *A. karschi*’s introduction to Pennsylvania will ever be determined. The species has a broad native range in East Asia extending from China and Taiwan to Japan (12), and it was recently reported in Korea (63). The Philadelphia region (including Delaware County) has had a 300 year history of trade with East Asia that may have included countless opportunities for accidental importation of a soil-associated spider among potted plants. Indeed, Nentwig (67) suggests that spiders introduced with potted plants have higher establishment rates relative to those introduced by other means. In the 1700s and 1800s Philadelphia was the center of American botany and horticulture and many plants from around the world, including Asia, were actively collected, imported and traded for exhibition and cultivation in public and private gardens (68,69). Many of the region’s great gardens and arboreta of that era still exist to some extent (70), including Tyler Arboretum (visited in this study) and Bartram’s Garden in west Philadelphia, the home of noted American botanists John Bartram and his son William (71,72). William Bartram’s contemporary in the late 1700s, William Hamilton, built his estate “The Woodlands” overlooking Philadelphia’s Schuylkill River and his gardens and greenhouse boasted of having every rare plant he’d ever heard of from around the world (73,74). In 1784, after the American Revolution, direct shipping trade began between Philadelphia and China and at its peak represented about a third of all US trade with China (75). A very significant Asian botanical importation event occurred later, in 1926, when the Japanese government presented 1,600 flowering trees to the City of Philadelphia to celebrate the 150^th^ anniversary of American independence (76). Regarding introductions of other soil-associated invertebrates, Asian jumping worms (*Amynthas* and *Metaphire* spp.) were presumably brought to the US in the 1800s in the soil of potted plants, and recent studies have shown that they displace native worms and are changing the soil where they occur (77). Coincidentally, nonnative jumping worms are present at many *A. karschi* sites in Pennsylvania (Tessler, personal observations).

### Conclusion

Many spider species have been accidentally introduced by humans to a new continent and became established (67), nearly all from the phylogenetically recent infraorder Araneomorphae. Within the more primitive mygalomorphs, the Mexican redrump tarantula (Theraphosidae) native to Mexico and Central America has become established in Florida USA (78). Presumably escaped from the pet trade, these tarantulas dig burrows and appear to be restricted to a small area with climate and habitat features similar to its native range. In this study we show that *Atypus snetsingeri* in Pennsylvania is genetically conspecific with *Atypus karschi* native to East Asia. The species appears to have been introduced by humans to Pennsylvania, probably in association with potted plants, and is now naturalized and locally common within a limited range that includes urban and forested areas. It is unlikely that the source or timing of the introduction can be determined in a region renowned for its colonial-era horticulturalists, elaborate international gardens, and long history of shipping trade with East Asia. This is the first case of an introduced species of Atypoidea from the infraorder Mygalomorphae.

## Author contributions

Performed the taxonomic and ecological observations: MR, ST. Performed the karyological analyses: IH, MF, JK. Performed the molecular analyses: PH. Prepared the figures: NG, ST, JK, PH. Wrote the paper: MR, ST, JK, PH, NG, IH, VR.

## Acknowledgements

We thank Tereza Kořínková for her comments on the manuscript, Marie L. Schmidt for her help with the research in Pennsylvania, and the Tyler Arboretum for permission to work on the property and collect specimens for scientific study. The study was supported by the Ministry of Education, Youth and Sports of the Czech Republic (project LTAUSA18171) and by the Ministry of Agriculture of the Czech Republic (project MZe RO0418). The chromosome part of this study was funded by the Charles University (project 92218) and Ministry of Education, Youth, and Sports of the Czech Republic (projects SVV-260568 and LTAUSA19142).

